# An epigenetic palimpsest: the King penguin’s blood methylome reveals conserved hematopoietic ageing

**DOI:** 10.64898/2026.10.10.757848

**Authors:** Mikaela Hukkanen, Flávia A. Nitta Fernandes, Leyla R. Davis, Maureen Hofer, Gaël Bardon, Elodie Paciello, Sandrine Zahn, Tianyu Zhu, Sofia Westerlund, Britta S. Meyer, Miina Ollikainen, Céline Le Bohec, Robin Cristofari

## Abstract

Age-associated DNA methylation is conserved across vertebrates, but whether epigenetic ageing reflects cellular ageing programmes or age-dependent shifts in cell composition remains unresolved. Here, using 126 whole-blood methylomes from known-age King penguins (*Aptenodytes patagonicus*), we exploit a vertebrate system in which nucleated erythrocytes dominate peripheral blood, minimizing the influence of leukocyte heterogeneity. We identify a core set of 409 ageing-associated genes showing lifelong, clock-like methylation change. These genes are strongly enriched for regulators of hematopoietic stem and progenitor cell (HSPC) function and niche communication. Integration of methylome, transcriptome and chromatin-accessibility data reveals that many of the strongest ageing signals derive from genes inactive in mature erythrocytes, consistent with the persistence of epigenetic states from ancestral hematopoietic progenitors. A comparison with methylation profiles from 1,501 humans identified a conserved cross-species subset of ageing genes, and a human epigenetic age model constructed solely from orthologues of penguin ageing genes predicts chronological age with high accuracy while remaining largely insensitive to blood cell composition. We propose that blood epigenetic ageing signatures contain a long-lived cellular memory of stem-cell ageing, that may represent an evolutionarily ancient layer of the vertebrate ageing process.

## Introduction

Age-associated DNA methylation changes are among the most reproducible molecular signatures of vertebrate ageing^1,2^. The general principle of epigenetic ageing appears robustly conserved in vertebrates: it is ubiquitous in mammals^3^, and appears to work in a broadly similar manner in birds^4,5^, frogs^6^, or teleost fish^7^, allowing for the development of CpG methylation-based age-predictors or “epigenetic clock” models in a wide array of taxa^3,4,6,8,9^. Yet the loci that underpin these clocks differ substantially between species: even between mammals, for example, methylation at the telomerase reverse-transcriptase promoter tracks age in the long-lived mole-rat, but not in the short-lived mouse^9^. Thus, the biological processes that drive the existence and remarkable reliability of epigenetic clocks remain poorly understood.

The most widely used epigenetic clocks are based on whole blood methylation: but how short-lived blood cells^10^ can reflect the age of the whole body remains controversial. Current interpretations are mainly structured around two non-mutually-exclusive hypotheses. On the one hand, the tissue composition hypothesis suggests that observed age-associated changes in methylation patterns mainly reflect changes in sampled cell subpopulations. This is supported by the observation that in humans, epigenetic clocks robustly reflect known, age-associated changes in blood cell counts^11^, and that different blood cell types exhibit different epigenetic ages^12^. On the other hand, the stem cell dynamics hypothesis proposes that most observed changes in methylation patterns hail back to changes happening in the stem cell compartment^1,13,14^. This hypothesis is mainly supported, so far, by mathematical models^14,15^, while empirical evidence remains ambiguous: after hematopoietic stem cell (HSC) transplantation, different studies have found that the epigenetic age of allogenic blood cells either conserved the age of the donor^16^ (supporting the stem-cell-age model) or drifted back towards the age of the host body^17^ (excluding a pure stem-cell-age model). Separately, injections of younger allogenic plasma rejuvenate blood epigenetic signatures in older rats^18^, suggesting that the blood methylome also responds to peripheral, non-stem signals. Thus, whether the ageing of the blood methylome reflects age-related changes of the stem cell lineage, changes in leucocyte composition, or a combination of both, is still an open question.

Non-mammalian vertebrates can help circumvent this dilemma. In non-mammalian vertebrates, the whole blood methylome is overwhelmingly dominated by nucleated red blood cells (RBCs, 99.5 to 99.9% of all blood cells^19^ in an adult bird). The expected contribution of leukocytes (0.1 to 0.5%) is thus of the same magnitude as the false-positive rate associated with cytosine methylation detection^20^, making the contribution of changes in immune cell composition minor to the whole blood DNA methylation landscape. This should, in theory, normalise the components of variance due to blood cell type changes, and either erase most age-associated methylation signals (if the tissue composition theory is sufficient), or reveal the underlying stem-cell-associated signal (if the stem cell theory holds across vertebrates).

Here, we test these two predictions by focusing on a long-lived bird model of ageing, the King penguin *Aptenodytes patagonicus*^5^ - a model that naturally resolves the cell-type-composition problem. Specifically, our aims are (1) to test for the persistence of clock-like epigenetic patterns in RBC-dominated data, (2) to assess whether, in the absence of confounding cell-type changes, we can detect a clearly stem-cell-associated signal, and (3) to evaluate whether the identified epigenetic ageing mechanisms are conserved across vertebrates. To achieve this, we combine a set of 126 whole blood methylomes, together with a reference panel of blood RNA and ATAC-seq, from known-age King penguins, to establish the core set of genes in the King penguin’s ageing methylome. As a human reference, we use methylation array data in ∼1,500 human twins. To formulate well-supported hypotheses as to the role of age-related DNA methylation change in penguin and human peripheral blood, we use published reference data for both mouse and human hematopoietic lineage cells. Overall, we show that RBC-dominated penguin blood retains a robust age-associated methylation signal that is clearly enriched for hematopoietic stem and progenitor cells (HSPC)-associated genes. This observation supports an evolutionarily conserved role of hematopoietic stem and progenitor cell dynamics to epigenetic ageing.

## Results

### Distinct signatures of time in the methylome

Time-associated changes in the King penguin’s methylome can be divided into (1) a large set of changes that happen in early life and stabilise at maturity, and (2) a much smaller set of truly ageing-associated changes that follow a consistent trajectory throughout life. Briefly, we produced genome-wide CpG methylation data by enzymatic conversion^20^ for 126 known age King penguin whole blood samples (ranging from 11 months to 35 years, see Supplementary Table 1). We first performed an epigenome-wide association study (EWAS) on ∼10 million filtered CpG sites, using logistic regression to model discrete 5mC counts as a function of *log*(age), sex, and technical covariates (see Methods). In a first approach, we selected the 47,384 CpG sites which carried the strongest age signal, with both a false-discovery-rate (FDR) corrected p-value ≤ 1e^-3^, and a change in |*log*(odds)| per unit of *log*(age)≥ 1.4 (in other words, an expected 4-fold change in methylation odds of each *e*-folding of age). Variation across these sites could be summarised as 5 typical age-related trajectories, inferred as clusters of CpGs with correlated methylation patterns using WGCNA^21^ (Fig. 1A). Trajectories of the types I, IV and V (see Fig. 1A) together accounted for the change observed in ∼70% CpG sites, and were all characterised by rapid positive (Type IV), negative (Type I) or sex-specific (Type V) change in early life (before age 2.5, the age of maturity in King penguins), followed by stabilisation or strongly attenuated change in adulthood. Types II and III, on the other hand, accounted for the remaining ∼30% sites, and were characterised by sustained lifelong clock-like change, either negative (Type II) or positive (Type III). Remarkably, while age-associated hypermethylation is an important part of the signal in humans and other mammals (>50% of Horvath’s clock-site set^22^, ∼40% of all age-associated 450k CpGs^23^, and in >90% of CpGs conserved across tissues in mammals^3^), hypomethylation is clearly the dominant signal in King penguin aging (92.4% of all sites are assigned to hypomethylated Types I, II or V).

**Fig. 1.**
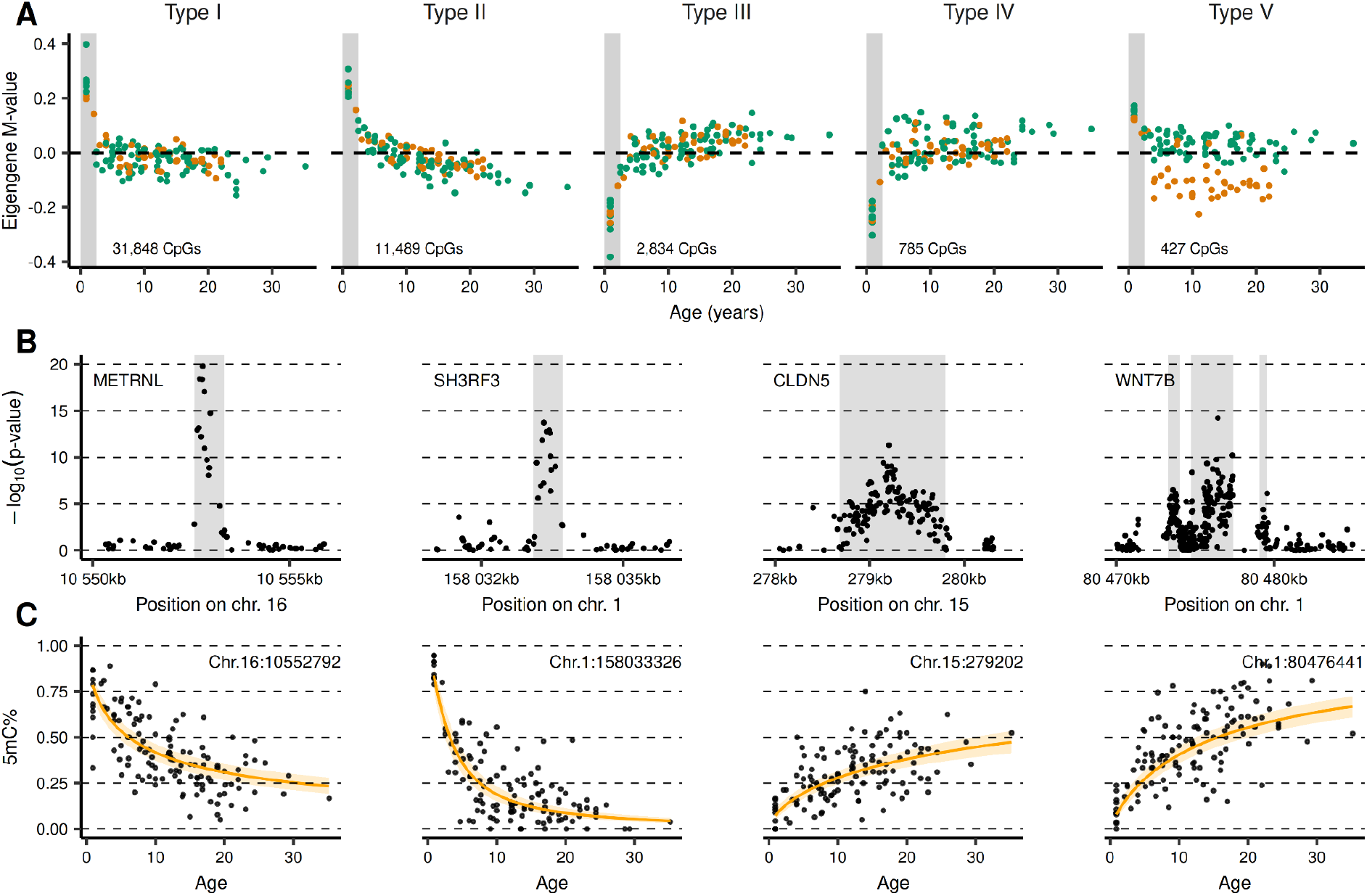
Time-associated signals are prevalent throughout the King penguin methylome. **(A)** Time-associated methylation change follows five types of trajectories (eigengenes), as inferred by WGCNA co-methylation clustering. In green, males, in orange, females. The gray area represents early-life (age < 2.5 years). The number of CpG loci assigned to each type is indicated in each panel, and panels are sorted by number of CpGs. Methylation level is normalised as M-value log_2_((C+ε)/(T+ε)) In trajectory types II and III, down- or up-methylation is consistent throughout life. In trajectory types I, IV and V, rapid change in early life is followed by weaker time-associated change. **(B-C)** The four strongest (lowest apex p-value) age-associated DMGs in the King penguin genome, removing early-life. METRNL and SH3RF3 follow a Type II trajectory, CLDN5 and WNT7B a Type III. **(B)** Local Manhattan plots for each of the four top aDMGs. The shaded gray area represents DMR coordinates. **(C)** Methylation probability as a function of age, at the apex site for each gene. Raw ratio of methylated by total cytosines (C/(C+T)). Orange line: fitted values predicted from our binomial models. Orange area: 95% prediction interval (mean ±1.96xSD from the model predictions).

### A restricted core set of ageing genes

We focused further on identifying specifically age-associated CpG sites. Our comethylation analysis suggested that early-life changes can drive most of the signal in e.g. Type I loci (Fig. 1A). Thus, we proceeded to identify age-associated differentially methylated regions (aDMRs) by limiting our EWAS analysis to adult birds (aged 2.5 to 35 years old, N=114). aDMRs were defined as clusters of significantly age-associated CpG sites after Stouffer–Liptak-Kechris correction for local autocorrelation^24^. We identified 578 aDMRs using the comb-p algorithm^24^ (counts reflect DMRs identified with a seed p-value≤1e^-4^, 500 bp windows, region Sidak-corrected p-value≤1e^-7^ - see Methods, and Supplementary Table 3 for the full list). Of these 578 aDMRs, 448 were associated with 409 unique age-associated differentially-methylated genes (aDMGs) (between 15kb upstream and 2kb downstream) - with some genes being associated with several DMRs (see Fig. 1B, WNT7B, for example). Within these aDMRs, CpG methylation follows a consistent and regular trajectory with log(age) (see Fig. 1C for illustration): 491 aDMRs are consistently hypomethylated with age, and 87 are hypermethylated with age. When tested against the King penguin-human orthology set (see Methods), these aDMGs are significantly enriched for transcription factors (48 genes, 1.6-fold enrichment, Fisher exact test FDR p-value 0.00104), protein kinases (27 genes, 2.3-fold enrichment, FDR 0.000289) and homeobox-domain genes (11 genes, 2.1-fold enrichment, FDR 0.0278). This suggests that methylation-controlled ageing genes have more regulatory than effector roles, and that ageing networks may be largely controlled from the top down through general regulatory switches.

### A signal carried by nucleated erythrocytes

Compared to humans, the set of King penguin age-associated differentially methylated sites (aDMS) covers a remarkably small proportion of all tested CpGs: 4,800 sites at FDR≤0.05 (0.049% of all tested sites), as opposed to the ∼40% to 50% of age-associated CpGs in human arrays^23,25^. Technical factors may contribute to that difference: our penguin sample size, in particular, is considerably lower. However, in our sensitivity analyses, human EWASes downsampled to an identical sample size had an average ∼3.7% of tested sites with FDR≤0.05, i.e. a >75-fold increase compared to the penguin EWAS. This suggests that limited sample size alone is unlikely to fully explain the difference (see Methods, Suppl. Fig. 1). The small proportion of aDMS in King penguins is also consistent with the expected reduction of cell-type composition components of ageing. Here, leukocytes contribute to only a minor fraction (<1%) of sequencing reads, while measured DNA methylation levels at age-associated apex sites vary between ∼100% and ∼0% (Fig. 1C): in other words, changes ultimately affect all sampled cells, necessarily including RBCs. Thus, the small number of penguin aDMS can be interpreted as reflecting (a) age-associated changes in yet-undocumented avian erythrocyte subtype proportions, and/or (b) inherited methylation profiles reflecting an ageing progenitor compartment. In any case, these patterns are practically unconfounded by variations in immune cells that only account for a negligible part of our data.

### Hematopoietic stem and progenitor cell biology dominates the ageing blood methylome

The ontology of the genes in our penguin aDMG set strongly favours the “progenitor methylation inheritance” hypothesis. First, we searched the transcriptomics-informed PanglaoDB marker gene database^26^ to identify enriched blood or bone cell type markers in our aDMGs: HSCs were the single most enriched cell type (3.3-fold enrichment, FDR-corrected p-value = 0.0285). We replicated this result independently using the literature-informed CellMarker2 database^27^, restricted to either human or mouse erythroid lineage cell types (see Methods). Against humans, markers for HSCs (Cell Ontology CL_0000037) were the only significantly enriched set (3.3-fold enrichment, FDR-corrected p-value = 0.0183). Against the mouse reference, the most enriched cell type was hematopoietic lineage restricted progenitors (CL_0002031, 5.96-fold enrichment, FDR-corrected p-value = 0.0491). This was followed by HSCs, although nominally-only significantly (CL_0000037, 1.5-fold enrichment, raw p-value = 0.0423, FDR *ns*). Our aDMGs are also enriched for genes that are expressed early during hematopoiesis, in mice. Briefly, we classified transcripts in the Haemopedia database28 according to their peak expression stage (see Fig. 2A), under the assumption that haematopoiesis is broadly conserved in vertebrates^29^. We tested whether our aDMGs differed from the full gene set in their distribution across hematopoietic stages by comparing observed stage counts to those expected under random sampling from the background set using Pearson’s chi-square statistic. Significance was assessed by 100,000 Monte Carlo simulations. We found that our aDMG set differed significantly from the complete penguin gene set p-value = 0.00127, see Fig. 2B-C), a signal driven largely by enrichment for genes peaking, in mice, at the multipotent progenitor stage, and by depletion for genes peaking at the erythroid-committed progenitor stages - see Fig. 2D.

**Fig. 2.**
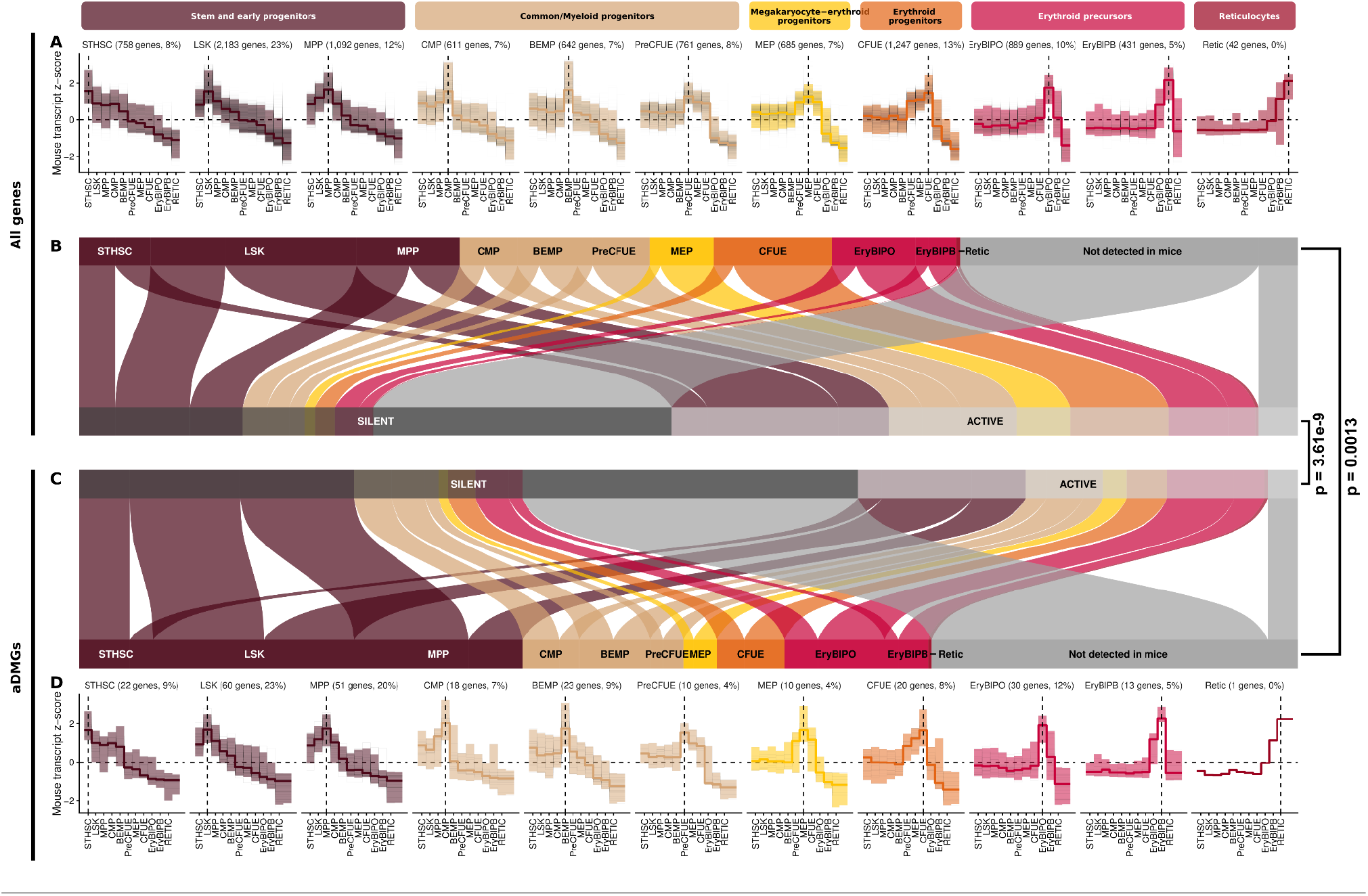
King penguin aDMGs are dominated by early-hematopoiesis genes. **(A)** and **(D)**: z-scores for gene expression along murine erythropoiesis (data from the Haemopedia mouse RNAseq dataset), classified by peak expression stage (STHSC: short-term hematopoietic stem cells, LSK: Lin(-)Sca1(+)Kit(+) cells, MPP: multipotent progenitors, CMP: common myeloid progenitors, BEMP: basophil/eosinophil/mast cells progenitor, PreCFUE: pre-colony forming unit-erythroid, MEP: Megakaryocyte-erythroid progenitors. CFUE: colony forming unit-erythroid, EryBlPO: proerythroblasts, EryBlPB: basophilic erythroblasts, Retic: reticulocytes), together with median and 95% interquantile range (coloured line and area). Colours refer to border stages of erythropoiesis (see legend at the top of the figure). The number of genes and percentage of the full gene set are given at the top of each subplot. **(A)**: full set of Haemopedia transcripts with a known ortholog in King penguins, and **(D)**: Haemopedia transcripts with an ortholog in the King penguin aDMGs. **(B)** and **(C)**: correspondence between each gene’s peak expression stage, and activity status in King penguin blood chromatin accessibility and gene expression data, in the full mouse-penguin orthology set **(B)** and the mouse-penguin aDMG orthology set **(B)**. In gray: most genes that are not detected in murine erythropoiesis are silent in King penguins. Right margin: p-values for statistical differences in gene activity status between aDMGs and the whole gene set (Fisher’s exact test), and in transcription peak stage distribution (exact permutation test with 1e^5^ replicates).

### HSC self-renewal and maintenance

Specifically, hematopoietic stem and progenitor (HSPC)-specific genes in our aDMG set included well-studied transcriptional regulators of HSPC identity, including MEIS1, MECOM, PRDM16, HOXB3-6, NFIA, RUNX2, JUNB or HLX. MEIS1 is one of the most consistently reported markers of long-term HSC establishment and quiescence^30,31^, including in birds^32^. It is hypermethylated during haematopoietic commitment^33^, but hypomethylated with age in our data (Sidak-corrected p-value = 4.061e^-40^), consistent with enhanced HSC self-renewal; its mechanistic partner PBX3^34^ is also an aDMG here. Other aDMGs with established roles in HSC quiescence or self-renewal include transcriptional regulator MECOM/EVI1^35,36^, metabolic regulator PRDM16^37^, HOXB3-6^38,39^, erythroid commitment regulator NFIA^40,41^, RUNX2^42^, JUNB^43,44^, CBFA2T3 and CBFA2T2^45^, SATB1^46^ (and its paralog SATB2^47^). Significantly, HLX must be repressed to maintain stemness in HSC^48^, and is one of the few genes that become hypermethylated with age in our aDMG set, in line with possible age-associated silencing in order to maintain stemness. On the other hand, several classical HSPC markers, such as KIT and HLF^49^, are absent from our aDMG set: however, while these are well-established stable markers for HSPCs, there is, to our knowledge, no evidence linking them to HSPC ageing in mammalian models either.

### HSPC and HSPC-niche

Several pathways classically associated with development are also clearly mobilised during the ageing process in our adult bird data. This includes, amongst others, Notch pathway members (NOTCH1, DLL4, HEY2, TCF3, LFNG) and non-canonical Wnt components (RYK, WNT3A, WNT7B, ROR2), but also a remarkably complete set of semaphorin receptor genes (PLXNA1, PLXNA2, PLXNA4, PLXNB2 and NRP2), and two ephrins (EPHA3 and EPHB1). Together with the HSPC niche regulator FOXC1^50^, with bone morphogenetic proteins BMP3 and BMP7, with caveolins CAVIN1 and 2, with as subunits α and β of the cholinergic receptor CHRNA3 and CHRNB4^51^, and with growth factors including PDGFRB^52^, EGFL7^53^, FGF1 and its receptor FGFR1, these pathways may be collectively involved in angiogenesis and the modelling of the bone marrow’s microenvironment. However, it is unclear how HSPC-niche methylation patterns could be transmitted into the differentiated blood lineage: instead, we propose that their differential methylation with age may be a sign of hemoatopoietic progenitor-to-niche signalling^54^ in the ageing process.

### General hallmarks of ageing in the blood compartment

King penguin blood epigenetic ageing also involves more “classical” hallmarks of ageing, such as the FOXO longevity signalling pathway^55^ (FOXO1) or the retinoid X nuclear receptor^56^ (RXRA), as well as genes relevant for autophagy regulation (e.g. WIPI1, RB1CC1, BCL2, or CTSB), or epigenetic remodelling (CHD6 and 7, SATB1 and 2, or PBRM1). At the same time, our aDMG set also includes genes that are clearly relevant to erythroid cells (and thus possibly to differentiated avian RBCs) including RBC surface antigen ACKR1/DARC, iron-reductase STEAP3, or heme transporter SLC48A1/HRG1. Changes in methylation at these loci could reflect either changes in RBC phenotype with age, or alternatively changes in the cytodemography of circulating RBCs, with the exact proportion of immature to mature cells changing slightly with age.

### NFI-A, RUNX2 and Jun-B transcription factor motif are enriched in aDMRs

So far, we showed that our aDMGs are enriched for upstream regulatory genes (including transcription factors and HOX genes), and for HSPC-associated genes. If these are indeed part of an HSPC-related regulatory network, we would also expect them to be associated with HSPC-related TF binding sites. To test this, we searched our aDMRs for known transcription factor (TF) binding motifs in the JASPAR core non-redundant vertebrate set^57^ (see Methods), and found significant enrichment for three HSPC-related TF classes. Specifically, out of the 277 enriched motifs, 3 classes including 37 individual motifs were both high-ranking and directly represented in our aDMG set. The first of these is the nuclear factor I (NF-I) TF class, including NFI-A (Bonferroni-corrected p-value = 7.03e^-27^, e-value = 1.58e^-23^), which is itself identified as an aDMG in this study (Sidak-corrected p-value = 6.195e^-09^). The second is the Runt domain class (RUNX), including RUNX2 (Bonferroni p-value = 3.23e^-16^, e-value = 7.27e^-16^), also an aDMG (Sidak-corrected p-value = 2.083e^-90^). The third is the Jun-B / activator protein I (AP-I) class, including JUNB (Bonferroni p-value = 4.71e^-14^, e-value = 1.06e^-10^), itself an aDMG (Sidak-corrected p-value = 2.470e^-26^). aDMRs are also strongly enriched for forkhead and homeobox motifs - see Supplementary Table 5 for the full list. To confirm this result, we also tested the aDMG against a database of externally-validated TF-gene associations in humans, using the MSigDB TFT gene set^58^. Consistently with our *de novo* search, top enriched gene sets included Jun-B / AP-1 (46 genes in overlap, FDR 1.39e^-15^), NF-I (33 genes in overlap, FDR 1.4e^-12^), RUNX (19 genes in overlap, FDR 1.59e^-10^ - see Supplementary Table 6). Overall, the consistent enrichment for HSPC-related TF binding motifs, together with the over-representation of HSPC-related genes in our aDMG set, supports the hypothesis that age-related methylation patterns observed in the King penguin’s erythrocytes reflect age-related epigenetic changes happening in, and inherited from the hematopoietic stem and progenitor compartments, and that are less influenced by changes in circulating blood cell composition.

### Core HSPC-linked aDMGs are not active in blood

To rule out the possibility that genes identified as HSPC-related in mammals are in fact expressed in differentiated blood in birds, we produced whole-blood RNA-seq and pure-type RBC ATAC-seq for 4 King penguins from our methylation population (see Methods). We found that genes with an open promoter were much more likely to be expressed than genes with a closed promoter (Fisher’s exact test: odds ratio = 27.3, 95% CI 25.0-29.9, p-value < 2.2e^-16^; n = 18,140 genes). Consistent with our hypothesis, we found that our aDMGS were significantly under-represented in active genes (i.e. at the same time open-promoter in RBC, and detectable in whole-blood transcriptomics - Fisher’s exact test: odds ratio = 1.28, 95% CI 1.04-1.59, p-value =0.0204) - see Fig. 2B-C. These include some of the most HSPC-relevant genes, including MEIS1, MECOM, PRDM16, or HLX (see Supplementary Table 4 for the full table). Symmetrically, an aDMG with clear relevance to mature RBC biology such as ACKR1/DARC, is both expressed in our data, and has an open promoter in RBCs, likely indicating a regulatory change in RBC methylation and expression through life. Importantly, genes that are not currently known to contribute to HSPC maintenance emerge in our data as possible candidates: in particular, the **Meteorin-like adipomyokine METRNL** is one of the clearest aDMGs (Sidak p-value 1e^-96^), but is not detectable in the blood transcriptome, and has a closed promoter area. METRNL was recently identified as a high affinity ligand for the KIT receptor (a canonical HSPC marker), and as a driver of angiogenesis^59^ - a process closely related to HSPC development. This makes METRNL a strong new candidate for HSPC maintenance in birds, and potentially in mammals.

### Conserved epigenetic aging signatures in King penguins and humans

The central role of nucleated RBCs in penguin epigenetic ageing raises the possibility that homology between avian and mammalian ageing is in fact rather low. To systematically evaluate this, we conducted an epigenome-wide association study (EWAS) using Illumina 450K BeadChip data from 1,501 human twins across 761 families (Supplementary Table 2). Out of 390,304 quality-controlled CpGs (annotated to 19,863 human genes), we focused on 11,380 human CpGs that directly mapped to 357 identified ortholog genes of King penguin aDMGs. After controlling for sex, lifestyle risk factors, and technical covariates, we found that human aging was significantly associated with methylation at 146of these CpGs, corresponding to 91 unique penguin aDMGs (FDR ≤ 1e^-3^; Suppl. Fig. 2). When correcting for changes in blood composition (12 white blood cell type deconvolution^60^), 77 CpGs spanning 60 distinct penguin aDMGs remained significantly age-associated. These aDMGs included established age-related genes included in pan-mammalian epigenetic clocks^3^ (e.g., PRDM16, FOXC1, TBR1, NFIA and MKLN1), genes identified in cell-type independent and mitotic human clocks (MGA in IntrinClock^61^; CHRNB4 and CCDC3 in stemTOC^62^), but also genes whose role in ageing is yet to be established (Fig. 3A). On the other hand, penguin aDMGs represent only a small subset (2.8%) of all human aDMGs, and several of the best-supported human aDMGs included CpG sites in our King penguin data, but were not detected as age-associated. For example, in our human EWAS, methylation at ELOVL2, FHL2, TRIM59, CCDC102B and KLHL35 genes is significantly associated with age, replicating previous findings on humans^63^, yet none map to penguin aDMGs. Conversely, several penguin aDMGs did not have any age-associated CpG sites in humans, including METRNL, HOXB3, HOXB5, PLXNA2, PLXNB2 and CASTOR2 all for which the median FDR adjusted p-value across all CpGs was above 0.70. Overall, our analysis suggests the existence of a small core of deeply-conserved aDMGs that replicate across penguins and humans, but also underlines the existence of original pathways uniquely involved in ageing in both branches.

**Fig. 3.**
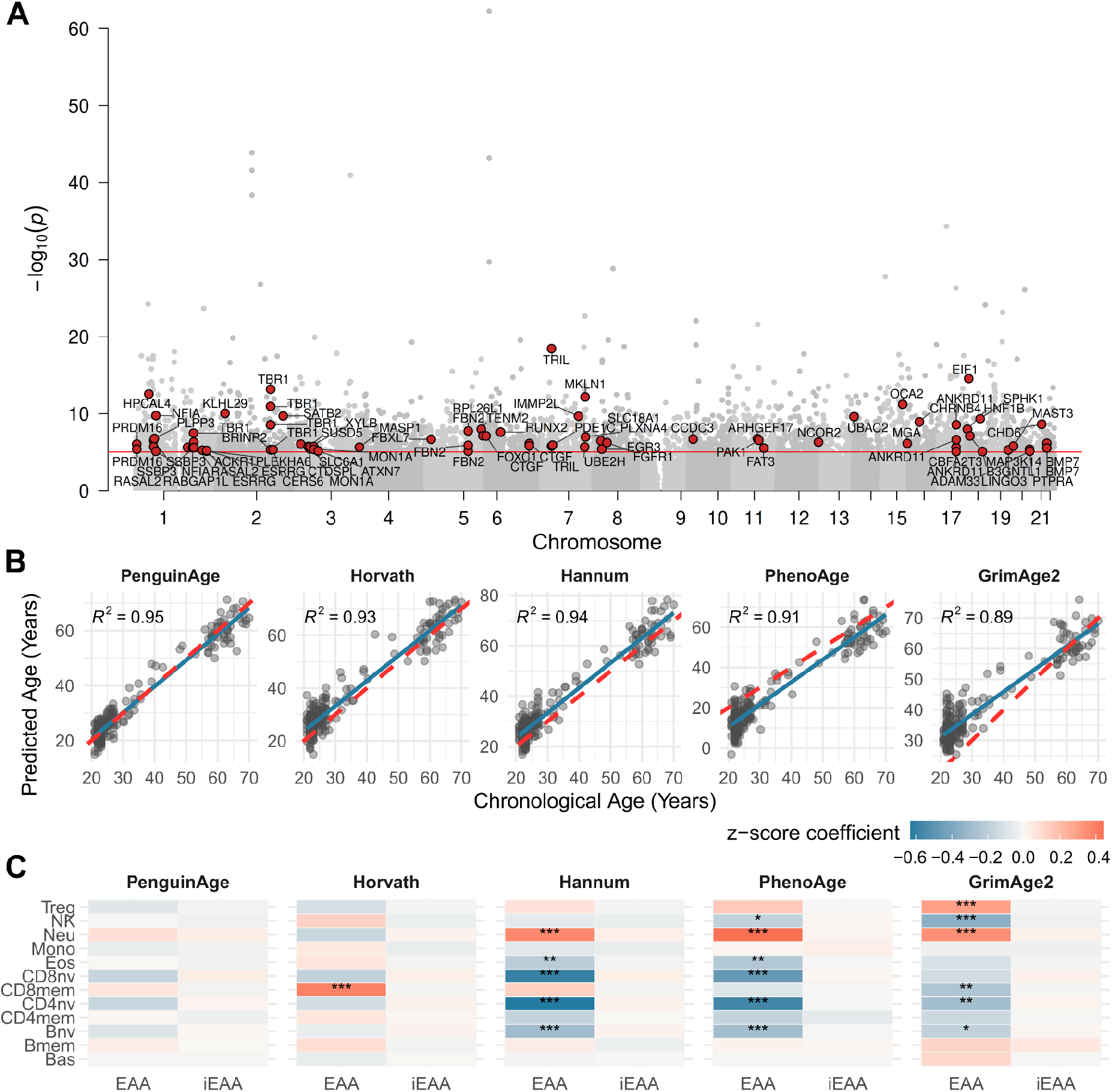
Epigenome-wide age association and epigenetic clock dynamics in humans. **(A)** Negative log_10_(p-values) for age are plotted against genomic coordinates (N_CpG_=390,304) from the epigenome-wide association study adjusted for cell-type proportions, BMI, smoking, alcohol, sex, technical factors, and family ID (N_Person_=1,501. N_Family_=761). The red line denotes the FDR = 0.001 threshold. Probes mapping to King Penguin aDMGs exceeding this threshold are highlighted in red. **(B)** Predicted versus observed age for the PenguinAge, Horvath, Hannum, PhenoAge and GrimAge2 epigenetic clocks (N_Person_=229). Solid blue lines show the linear regression; red dashed lines illustrate the 1:1 identity line. **(C)** Association of epigenetic age acceleration (EAA) and cell–type adjusted EAA (iEAA) with 12 cell types, adjusted for sex, age and age^2^. Tile color indicates model z-scores for each cell type; asterisks denote statistical significance based on FDR-adjusted p-values (***p < 0.001, **p < 0.01, *p < 0.05).

### A penguin-informed human clock predicts human age independently from cell type

To test whether King penguin aDMGs indeed represent an evolutionarily conserved, blood-cell-type-independent subset, we built an epigenetic age-prediction model (PenguinAge) trained on the subset of human CpGs mapping to penguin aDMG orthologs. Consistently with our hypothesis, this model demonstrated very robust, and largely cell-type-independent chronological age prediction in humans. To build this trans-species model, we extracted a subset of human 450K probes mapping to penguin aDMGs within 2kb of the aDMR (N_CpGs_=10,640, N_Genes_=327) across family-unrelated singletons (N=761). Using a 70:30 dataset partition, an elastic net regression model with 10-fold cross-validation was optimized on the training data to select the most predictive epigenetic features (330 CpG sites across 173 genes). The finalized PenguinAge clock displayed strong predictive performance, achieving a high-precision fit in the training set (R^2^ = 0.99, MAE = 1.22 years, N = 532) that satisfactorily generalized to the independent testing set (R^2^ = 0.95, MAE = 2.72 years, N = 229).

To evaluate PenguinAge against conventional human frameworks, we cross-benchmarked its performance in the testing dataset alongside four established epigenetic clocks. These clocks included the Horvath^22^ and Hannum^64^ first-generation clocks, which were trained directly on chronological age. We also compared it to second-generation clocks PhenoAge^65^ and GrimAge2^66^, which were trained on physiological risk phenotypes, lifestyle metrics (like smoking pack-years), and multi-morbidity mortality data to capture “biological age” and disease risk. The highest predictive power was observed for PenguinAge, followed by Hannum, Horvath, PhenoAge, and GrimAge2 clocks, with R^2^ ranging from 0.89 to 0.95 (Fig. 3B). Furthermore, PenguinAge’s predictive accuracy exceeded that of 998 out of 1,000 of clocks trained on randomly sampled human-penguin orthologue gene sets of the same size (see Methods), suggesting that its superior performance is specific to the penguin aDMGs rather than a combination of any random genes (Suppl. Fig. 3A).

We investigated the confounding impact of immune cell composition on clock metrics, reproducing the approach by Zhang and colleagues (2024)^11^. We modeled the associations of both raw Epigenetic Age Acceleration (EAA) and 11-cell-type-adjusted intrinsic EAA (iEAA) against 12 leukocyte subsets^11^ (Fig. 3C). Standardized multivariable regression controlling for sex, age, and quadratic age effects (Age^2^) revealed that, while many first- and second-generation human clocks remained sensitive to variation in immune cell composition, the PenguinAge framework predicted age independently from blood cell type composition. Moreover, PenguinAge showed weaker overall associations with blood cell composition than 96.7% of random gene clocks (Methods, Suppl Fig. 3B-C), further indicating that its robustness to cell composition is a distinctive feature of the penguin aDMGs underlying the clock. Interestingly, among the published clocks tested here, only the Horvath clock EAA metrics also showed minimal associations with cell types, only associating with memory B lymphocytes. This might be related to the multitissue training of the Horvath clock in comparison to the Hannum, PhenoAge and GrimAge2 clocks that have been trained on blood methylation only^22^. The fact that the PenguinAge clock appears to “see through” the variations in blood cell composition supports the idea that differential regulation of penguin aDMGs happens not in the blood itself, but upstream of blood cell differentiation, possibly in the HSPC compartment that is ancestral to all blood cells, in penguins as in humans.

## Discussion

Epigenetic clocks are now a well-established hallmark of vertebrate ageing, yet the cellular origins of their signal remain elusive^15,67,68^. By taking advantage of a long-lived avian system where nucleated erythrocytes constitute virtually the entire sequenced cellular fraction of blood^19^, we decoupled blood epigenetic ageing from the confounding effects of leukocyte composition inherent to mammalian blood^11^. This resulted in a considerably reduced set of aDMGs, compared to mammals. On the other hand, the resulting core set of age-associated differentially methylated genes was consistently linked to regulators of HSPC maintenance, self-renewal and differentiation, including MEIS1, MECOM, PRDM16, NFIA and members of the HOXB cluster. Taken together our results show that while cell-type composition is indeed a major driver of mammalian epigenetic clocks, another consistent component of the blood age-related signal is a cell-independent, evolutionarily-conserved legacy from the HSPC compartment, inherited by differentiated descendant cells. Importantly, these two views are complementary: age-related changes in HSPC contribute to immunosenescence^69^, and thus the observed changes in circulating blood cell types can themselves be interpreted as a result of age related changes in HSPC.

### The palimpsest in the ageing methylome

The observation that many of the strongest age-associated genes are apparently inactive in mature erythrocytes suggests a broader interpretation of blood epigenetic ageing. Several prominent aDMGs (including MECOM, PRDM16, FOXC1 and multiple HOXB genes) showed closed promoter chromatin and no detectable expression in circulating blood cells at all ages, yet retained some of the strongest age-associated methylation signals. This decoupling of gene activity from methylation signals in mature erythrocytes supports what we describe as an epigenetic palimpsest model: differentiated blood cells retain an inherited, archival imprint of epigenetic remodelling that occurred within the ancestral HSPC compartment. During differentiation, specialized terminal programs are written over, but ancestral stem cell epimutations remain decipherable at silenced loci, propagating downstream as a stable biological record. In line with this hypothesis, the genes responsible for maintaining or re-establishing DNA methylation profiles in successive cell division, TET and DNMT3, are the most frequently mutated genes in age-related clonal hematopoiesis^70,71^, which underscores the importance of methylome information conservation and modification through life for healthy hematopoiesis.

This palimpsest model provides a clue as to why cross-species epigenetic clocks generalize across divergent lineages^3^ despite differences in blood composition. In humans, epigenetic clocks have often been criticized for being confounded by age-associated shifts in white blood cell proportions^11,60^. However, our trans-species PenguinAge model, trained in humans and strictly on human orthologues of penguin aDMGs, predicted chronological age while remaining largely insensitive to leukocyte counts. The retention of these orthologous signals across enucleated-RBC (mammalian) and nucleated-RBC (avian) lineages demonstrates that epigenetic clocks are not pure downstream reflections of immunosenescence. In addition to revealing the deep evolutionary roots of epigenetic ageing, these conserved stem-cell dynamics offer a powerful avenue to refine biological clocks that directly report on stem cell viability, a process that has widespread effects on organismal health^71^.

### Ageing in HSPCs and their niche

This interpretation is also consistent with current views of hematopoietic ageing. Ageing HSPCs suffer from reduced regenerative capacity, increased clonal hematopoiesis^71,72^ and progressive loss of lymphoid potential in parallel with higher proportion of HSPC clones contributing to myelopoiesis^73,74^. Our results cannot distinguish between cell-lineage intrinsic effects, and changes in the relative contribution of distinct HSPC clones^71^: but both hypotheses predict that age-related changes within the stem-cell pool should become detectable in downstream blood cells. In this context, the enrichment of genes involved in stem-cell maintenance versus proliferation indicates that age-related methylation patterns record a fundamental fitness trade-off within the HSPC pool, shifting between high-output proliferation and maintenance-oriented clones. Consistent with this idea, lymphoid-committed progenitors have a younger epigenetic age^75^ than both differentiated bone marrow cells and their chronological age, and thus the reduced contribution of this lineage might contribute to increased epigenetic ageing with time, indicating that epigenetic clocks are sensitive to stem-cell state and differentiation history.

Beyond intrinsic stem-cell biology, our data also point towards age-associated remodelling of the stem-cell niche. Notch, Wnt, semaphorin and ephrin signalling components were strongly represented among aDMGs, together with established regulators of HSPC-microenvironment interactions such as FOXC1, EGFL7 and PDGFRB. These pathways coordinate vascular interactions and niche organization in vertebrate bone marrow and suggest that the ageing signal captured in blood reflects not only the state of HSPCs themselves but also their interaction with supporting cellular environments. More generally, this observation is consistent with emerging views that stem-cell ageing is strongly shaped by local microenvironments and intercellular communication^76–78^ rather than solely by cell-intrinsic events.

### The other layers of the palimpsest

Importantly, we do not suggest that the erythrocyte methylome is exclusively a readout of stem-cell ageing. In birds, specific aDMGs likely represent erythroid-specific biology, as illustrated by genes such as ACKR1/DARC, STEAP3 and SLC48A1. At the same time, the more than ten-fold shrinkage in the number of aDMGs between mammals and penguins is, by itself, a strong indication that circulating blood cell counts are major contributors to observed age-associated epigenetic changes in blood. Finally, additional components may arise from systemic physiological signals in circulating blood^17,79^, epimutational drift^71^, or broader organismal ageing processes that indirectly influence blood production and composition (Figure 4). We therefore view the ageing methylome as a layered structure in which multiple biological processes leave superimposed signatures. The key contribution of the king penguin model is that it largely removes the masking effects of leukocyte heterogeneity and reveals with unusual legibility a long-lived stem-cell-associated component of this epigenetic palimpsest.

**Fig. 4.**
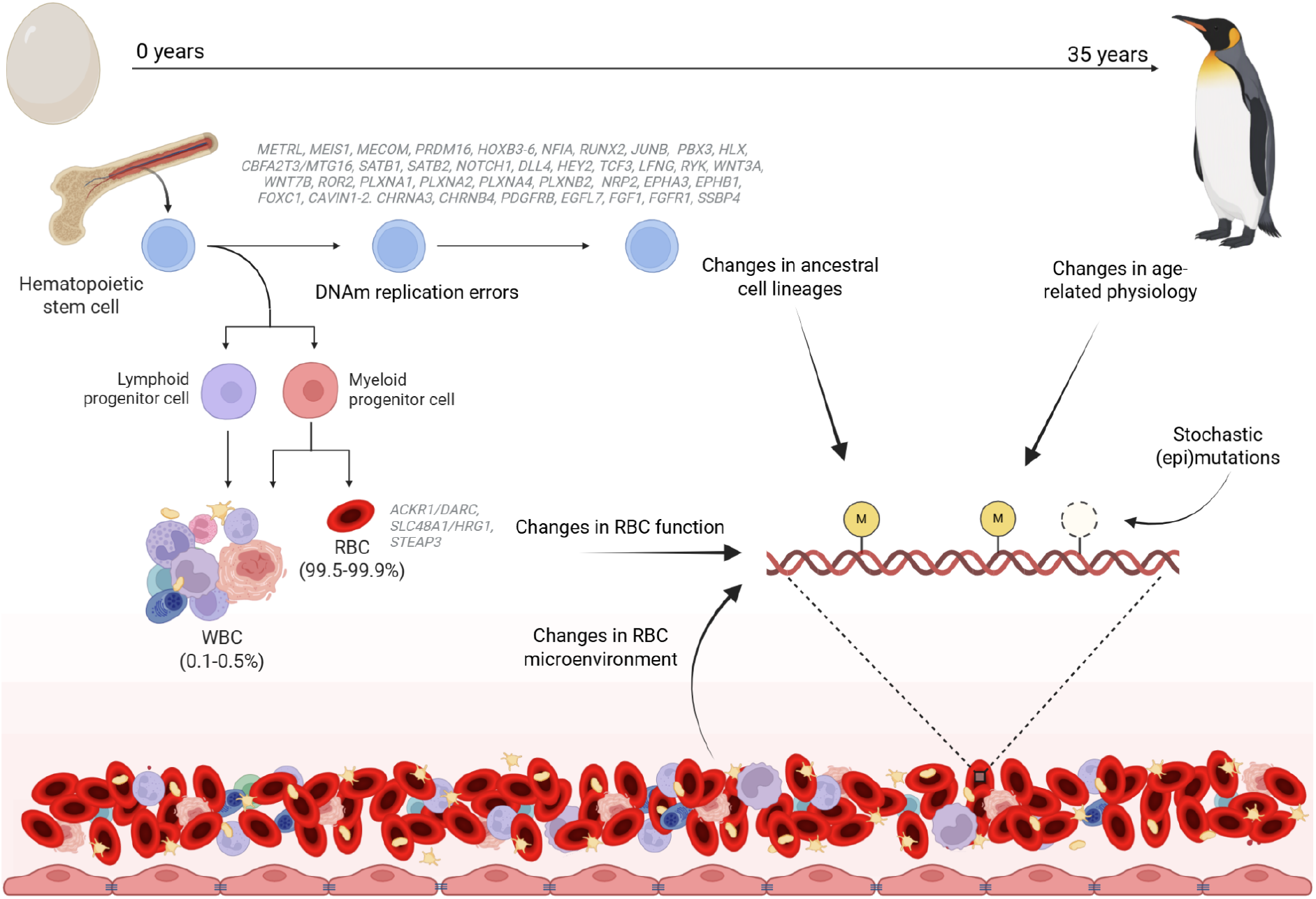
The palimpsest of the ageing methylome. Lifelong replication and ageing of hematopoietic stem cells generate methylation changes that are propagated through descendant blood lineages and captured in nucleated erythrocytes. Additional contributions arise from erythrocyte-specific biology, age-related changes in the blood microenvironment and organismal physiology, and stochastic epimutational drift. Created in BioRender. Hukkanen, M. (2026) https://BioRender.com/c44k91v.

### Limitations of this study

An important limitation of our study is that our data was restricted to peripheral blood DNA methylation without paired multi-omic or clinical health indices. While our primary objective was to compare avian and human epigenetic aging, such metrics would have strengthened our functional inferences. Second, we lacked an independent validation cohort of known-age king penguins, and while the human twin cohort served as a cross-species test, we could not validate the PenguinAge clock across distinct human populations. Third, cell-type enrichment models and functional annotations were informed by mammalian databases under the assumption of core hematopoiesis conservation, as comprehensive avian-specific functional annotations remain limited. Finally, cross-species translation was constrained by the incomplete genomic coverage of the Illumina 450K BeadChip, lacking probes for key penguin loci (such as the second intron of METRNL, identified as a major King penguin aDMR). Probe coverage limitations likely masked genuine orthologous aging signals in humans: in several cases, penguin aDMGs did not correspond to significant aDMS in humans, but are nevertheless validated components of established pan-mammalian epigenetic clocks and eutherian blood EWAS^3^. The limited comparative equivalence of our whole methylome aDMG set and the 450K chip highlights the need for large scale methylation analyses across taxa. Future replication in denser microarrays or using whole-methylome sequencing in other vertebrates may shed additional light on the role of individual genes in conserved vertebrate ageing.

### Conclusion

Overall, the King penguin system demonstrates that vertebrate blood epigenetic clocks are not a pure byproduct of age-associated shifts in leukocyte proportions, but also reflects a clear and evolutionarily-conserved signal inherited from HPSC aging. We show that differentiated nucleated erythrocytes retain in their methylome a palimpsest of stem-cell epimutations: at the same time, our results also strongly suggest that a similar palimpsest is still also decipherable in the human leukocyte methylome, albeit far more heavily over-written. Finally, we illustrate the usefulness of this signal as a peripheral biomarker by showing that orthologs of avian erythrocyte-aging loci can inform a high-precision, leukocyte-insensitive human epigenetic clock. We suggest that such a clock, by targeting specifically a known HPSC-aging-associated signal, can be applied to monitor typical age-associated ailments such as stem-cell pool exhaustion, immunosenescence, and clonal hematopoiesis risk.

## Methods

### Methylation data production

Sampling complied with the relevant ethical regulations. 126 DNA samples were collected from the whole peripheral blood of 116 King penguins of known age and sex, with repeated longitudinal sampling for 7 of them (between 2 and 3 samples for each of these, separated by at least 5 years, see Supplementary Table 1). 64 of these samples were re-used from a previous study^5^ (SRA accession number PRJNA1187342). Samples were collected either in Possession Island, Crozet Archipelago (N = 88) or from European zoos (N = 38). In the wild, the sampling was approved by the French ethics committee (latest: APAFIS#29338-2020070210516365) and the French Polar Environment Committee, and permits to handle the animals and access the breeding sites were issued by the “Terres Australes et Antarctiques Françaises”. In zoos, samples were collected either as part of routine veterinary examination, or approved by the Cantonal Veterinary Office of Zurich (license no. ZH033/2024; national reference no. 36243). After collection, blood was immediately frozen either with no buffer or in Queen Lysis Buffer solution and DNA extractions were performed with the NucleoSpin Blood Kit following the manufacturer’s protocol (Macherey-Nagel, Duren, German).

Whole-genome enzymatic-conversion sequencing libraries were prepared using the NEB EM-seq kit^20^, and sequenced on the Illumina NovaSeq 6000 platform either at CCGA in Kiel, Germany, or with GeneWiz (Azenta). Raw reads were trimmed, removing leading and trailing bases if the quality score dropped below 3 and retaining only reads of length > 36 bp, using Trimmomatic^80^ (v0.39). Reads were aligned to the King penguin’s reference genome GCF_965638725.1^81^ using bsbolt^82^ v1.5.0, and deduplicated and filtered using samtools^83^ v1.16.1. The reference genome was refined using (1) gCluster^84^ (GitHub commit version #248b42d) to identify CpG islands, (2) GUSHR (https://github.com/Gaius-Augustus/GUSHR, GitHub commit version #ee26d5c) and whole-transcriptome data for the King penguin^85^ aligned to the King penguin genome using HiSAT2^86^ v2.2.1, to annotate untranslated regions, and (3) PANNZER^87^ to add annotation to uncharacterised genes based on protein function prediction.

Individual methylated cytosines were called using bsbolt^82^. We calculated per-CpG average cytosine methylation level by pooling C and T calls from both strands at each CpG dinucleotide. We masked from our dataset all known C/T (or G/A) SNP positions based on a curated database of 40 King penguin whole genomes. Mean conversion efficiency, as assessed through Lambda phage unmethylated control false-positive call rate, was high at 99.7% (range: 96.4% to 99.8%). Finally, CpG sites were filtered to keep CpG loci with an average depth between 8 and 40x across all samples, an individual depth between 4 and 80x per individual, a minimum median absolute deviation per a site of 0.02, and a maximum of 40% missing data, for a total of ∼10 millions (9,933,293) CpG sites. More stringent filtering was applied for additional analyses; it is described below where relevant.

### RNA-seq and ATAC-seq

We produced whole-blood RNA sequencing and pure-type RBC ATAC-seq data for 4 King penguins sampled at Zoo Zürich under approval by the Cantonal Veterinary Office of Zurich (license no. ZH033/2024; national reference no. 36243), including 2 males and 2 females (see Supplementary Table 1).

For RNA-seq, 50µL whole blood were stored in RNAShield buffer (Zymo Research) and frozen at −80°C until RNA was extracted using the Zymo Blood RNA kit. Total RNA sequencing was performed after rRNA and globin-RNA depletion by Genewiz (Leipzig, Germany). Raw reads were trimmed^80^ and aligned to the King penguin reference genome using STAR^88^ (v2.7.11b), and transcript counts were calculated using HTSeq^89^ (v2.0.3). Counts were analysed in R using DESeq2^90^ (v1.44.0). Conservatively, a gene was considered not expressed in blood if (i) less than half the samples had a raw count ≥ 10, and (ii) the gene had a summed raw count across all 4 samples ≤ 40. Here, a total of 10,777 genes were found to be expressed in whole blood, and 7,363 showed no expression beyond noise level. RBCs dominated the signal, with the erythrocyte ankyrin ANK1 and the band 4.1 EPB41 transcripts among the 10 most expressed genes in our data, and ferritin heavy chain 1 (FTH1) being the single most expressed gene, all three with expression in the order of 5-10,000 CPM (mean raw count in the order of 10^5^). Immune cells are also detectable contributors, with e.g. leukocyte common antigen PTPRC in the order of 100 CPM.

For ATAC-seq, 300µL were resuspended in 600µL PBS, layered over 900µL Histopaque^®^-1119 (1.119 g/mL), and centrifuged at 400xg for 30 minutes at room temperature. The RBC phase was resuspended in 600µL PBS, and centrifuged again at 400xg for 15 minutes. The pellet was resuspended in 1800µL PBS. Purity of isolated RBCs was assessed visually in a Neubauer cell. Finally, 4µL of that suspension the cells were resuspended in RPMI medium containing 40% heat-inactivated FBS and 10% DMSO, and frozen at −1°C/minute to −80°C. ATAC-seq library preparation and sequencing were performed by Genewiz (Leipzig, Germany). Raw reads were trimmed^80^ and aligned using bowtie2^91^ (v2.5.3). After cleaning, peaks were called using MACS3^92^ (v3.0.1), with a q-value cutoff at 0.05. Signal strength was good, with fractions of reads in peaks > 0.35 in all four samples. Between-sample repeatability was also high, with all pairwise Jaccard indices > 0.65. Promoter openness was defined as the presence of an ATAC-seq peak within 500bp of the TSS position. In pure-type RBCs, 8,847 genes had an open promoter in at least one sample, and 7,660 an open promoter in all 4 samples. For analysis, we considered a gene as “open-promoter” if it had an open promoter in at least 3 out of 4 samples.

### Epigenome-wide association patterns

An epigenome-wide association study (EWAS) was performed on the 9,933,293 CpG sites that passed filtering. At each CpG site, a separate binomial model was fit using glmmTMB^93^ v1.1.11 in R, with the number of methylated and unmethylated reads at each position as a response, and including in all cases origin (“zoo” or “wild”), centered-scaled library conversion efficiency (as estimated using fully un-methylated Lambda control DNA^20^), centered-scaled mean genome-wide CpG methylation, and individual identity as a random effect to account for longitudinal sampling in a subset of birds (see Supplementary Table 1). We first modelled individual CpG methylation probability as a function of a logarithmic transform of age, including sex as an additive two-level covariate. Then, in order to exclude the effects of growth and to identify specifically ageing-associated loci, we restricted our dataset to birds aged 2.5 and above (when King penguins reach sexual maturity and become visually indistinguishable from adults). On the other hand, to identify loci where methylation changes sharply during early life, but that are not necessarily ageing-related (e.g. linked to the activation of reproductive programs, and the deactivation of initial growth pathways), we modelled age discrete, 2-level factor with a break point at 2.5 years of age. We considered a possible interaction with sex (to account for sex-specific changes during sexual maturation). In that case, we used emmeans^94^ to assess effect size and statistical significance.

### Co-methylation network analysis

In order to evaluate general patterns in DNA methylation change with age, we built a co-methylation network using WGCNA^21^. Based on our initial EWAS results, we first identified all CpG sites where DNA methylation broadly correlated with age across the whole lifespan, including log(age) as a linear predictor from hatching to old age, and sex as an additive covariate. Every CpG site with an FDR-corrected p-value ≤ 1e^-3^ and a log(odds) value for log(age)-related change in methylation probability was included in the analysis (47,383 CpGs). Methylation level at these sites was converted to M-value as log_2_((nC+ε)/(nT+ε)), with ε=1 (one pseudo-read to avoid division-by-zero issues) in order to normalise distributions^95^. Scale-free topology was reached with an exponent of 7 in our network. The topological overlap matrix was clustered using hierarchical clustering, and modules were identified using WGCNA’s cutreeDynamic function, with a splitting depth parameter set to 4 (very sensitive) and a minimum module size of 200 CpGs. Similar modules were finally re-merged whenever their eigengene correlation exceeded 0.1.

### EWAS peak identification

Since whole-genome CpG methylation signal is highly autocorrelated (as opposed to e.g. array data), classical false-discovery-rate correction assumptions do not hold. Here, we used *comb-p*^24^ (github commit #c18e137) to account for autocorrelation along neighbouring CpG sites. Peaks were nucleated with a raw EWAS p-value ≤ 1e^-4^ and CpGs were considered within a distance threshold of 500bp. We retained peaks with a combined and Šidák-corrected p-value ≤ 1e^-7^, on the condition that they contained at least 3 CpGs. These age-associated differentially methylated regions (aDMRs) were to their nearest gene using bedtools^96^ and custom bash/awk scripts. When an aDMR overlapped with several features, features were attributed according to the following ranking: first exon, other exon, 5’-UTR, first intron, other intron, 3’-UTR. A gene was considered possibly associated with an aDMR if the aDMR was located at a maximum of 15kb upstream of the gene or 2kb downstream of the gene. We also considered a narrower frame, from 2kb upstream to 2kb downstream of each gene. The genes associated with aDMRs are considered candidate age-associated differentially methyled genes (aDMGs).

### Gene set enrichment analysis

Gene set enrichment analysis in King penguins raises important questions regarding background reference gene sets. Here, we restricted all analyses to the set of genes for which orthology relationships were clearly established in the reference datasets. Gene symbols were normalised when necessary using HGNChelper^97^. For gene family enrichment annotations, we used the MSigDB database^58^, excluding cancer-related families to reflect our model system. For cell type enrichment, we used two independent databases. First, we used the curated single-cell gene expression database the PanglaoDB^26^, restricted to blood and bone tissues. Second, we used the CellMarker 2.0 literature-based database^27^, restricted to cell types within the broader erythroid lineage, i.e. Cell Ontology CL_0000037 (hematopoietic stem cells, HSC), CL_0000038 (erythroid progenitors, EP), CL_0000049 (common myeloid progenitors, CMP), CL_0000050 (megakaryocyte erythrocyte progenitors, MEP), CL_0000232 (erythroid cells),CL_0000763 (myeloid cells), CL_0000765 (erythroblasts, EryB), CL_0000837 (multipotent progenitors, MPP), CL_0000839 (common myeloid progenitors, CMP), CL_0000988 (hematopoietic lineage restricted progenitors), CL_0002031 (hematopoietic progenitors), and CL_0002034 (long-term hematopoietic stem cells, LT-HSC). For this literature-derived search, we analysed separately human and mouse datasets. We used Fisher’s one-sided test in R (version 4.3.3) to assess significance of overlap, correcting p-values for false-discovery.

For identifying the cell types expressing our aDMGs, we built a reference dataset based on the Haemopedia mouse haematopoietic data^28^. We downloaded raw count data, and selected the subset of cells that may be ancestral to circulating RBCs (the source of our DNA methylation data), namely, in order of differentiation: short-term haematopoietic stem cells (STHSC), Lineage Negative Sca+ Kit+ Multipotential progenitor (LSK), Multipotential progenitors (MPP), Common myeloid progenitors (CMP), Bipotential erythroid megakaryocyte progenitors (BEMP), Pre-erythroid colony forming units (PreCFUE), Megakaryocyte erythrocyte progenitors (MEP), Erythroid colony forming units (CFUE), Polychromatic erythroblasts (mixed with basophilic: EryBIPB; and mixed with orthochromatic: EryBIPO), and reticulocytes (Retic). Raw counts were normalised to transcripts-per-million. We restricted our analysis to genes that had a well-established orthology relationship between mice and King penguins (12,874 protein-coding genes). We used WGCNA^21^ to identify “typical” gene expression patterns throughout erythroid commitment (briefly, we used the blockwiseModules function, with an exponent of 2, a minimum module size of 15, and default sensitivity parameters to define modules of co-expression). Module assignments for the whole penguin-mouse orthology set, and for the aDMG-mouse orthology set, were compared using a chi-squared test with 6 degrees of freedom. Differences in individual modules were assessed based on chi-squared residuals, which were treated as approximate Z-values and converted to p-values.

### Motif discovery and transcription factor binding site enrichment

We used AME^98^ to test for known transcription factor (TF) binding site enrichment in the aDMRs. For short aDMRs (<300bp), the region was extended up- and down-stream up to 300bp. We generated a reference background using the sliding-window approach implemented in biasaway^99^, that matches local variation in sequence composition, from the King penguin reference genome masked for repetitive elements^81^ and for our aDMRs (to avoid data leakage), using the JASPAR 2026 CORE vertebrate non-redundant^57^ TF database. We used an adjusted p-value threshold ≤ 0.05 and an e-value ≤ 10. Analysis was performed on the MEME suite server^100^. Independently, we searched the MSigDB Transcription Factor Targets (TFT) gene sets using GSEA^58^. The TFT collection is based on TF motif enrichment in model species, and is entirely independent from our AME motif search in the King penguins: thus, both approaches represent essentially separate lines of evidence.

### Human EWAS

The human sample set used in this study includes 1,501 participants from 761 families of the population-based Finnish Twin Cohort (FTC)^101^. The FTC study has been approved by the appropriate national research ethics committees, with the most recent approval granted by the Hospital District of Helsinki and Uusimaa ethics board in 2018 (#1799/2017). Blood samples for DNA analyses were collected from each participant after they had signed a written informed consent. The samples and data used for the research were obtained as part of the Finnish Twin Cohort studies and have been deposited in the Biobank of the Finnish Institute for Health and Welfare in accordance with the Finnish Biobank Act (2013). The authors state that all procedures involved in this study adhere to the ethical standards of human experimentation, as well as to the principles of the Declaration of Helsinki.

Peripheral blood samples for the cohort participants were taken at an age ranging from 21 to 72 years (Supplementary Table 2). Lifestyle-related information for the participants was obtained from questionnaire data at the time of blood sampling^101^. Genome-wide DNA methylation was assessed using the Illumina Infinium HumanMethylation450 BeadChip, and raw intensity data were preprocessed and normalized as described in Hukkanen et al. (2026)^102^. Beta-values (proportion on methylation ranging between 0-1) were log2-logit transformed into M-values for statistical analyses.

To evaluate associations of M-values with chronological age while controlling for twin clustering, we used Generalized Estimating Equations (GEE). GEE models were computed in R (version 4.3.3) using the package gee version 4.13-29 with an exchangeable correlation structure. Both models were adjusted for technical covariates (BeadChip row and date, and sample-specific, z-score-scaled mean array-wide M-value), sex, and lifestyle-related covariates. These included BMI (kg/m^2^) as centered linear and quadratic terms, smoking status (current, former, never) and alcohol consumption (abstainer, low (<25 g per day), medium (<45 g per day), high (≥45 g per day) as further described in Zhao et al. (2023)^103^; Supplementary Table 2). The second model included additional covariates for relative cell-type proportions for 11 leukocytes including basophils (Bas), neutrophils (Neu), naive B-cells (Bnv), memory CD4+ T-cells (CD4mem), naive CD4+ T-cells (CD4nv), memory CD8+ T-cells (CD8mem), naive CD8+ T-cells (CD8nv), eosinophils (Eos), monocytes (Mono), natural killer cells (NK), and regulatory T-cells (Treg) computed via FlowSorted.BloodExtended.EPIC^60^. Memory B-cells (Bmem) were excluded as the baseline reference group due to having the lowest average population proportion across cell types with <1% of 0 proportions, following the logic described in Zhang et al. 2024^11^. Robust sandwich estimators were applied to compute Wald test p-values. Multiple testing corrections were performed using the Benjamini-Hochberg False Discovery Rate (FDR), with statistical significance defined at FDR<0.001.

Genomic features were annotated using the Illumina manifest B5 panel with UCSC RefGene database. To identify shared cross-species aging signals, we restricted the above-described King penguin age-associated differentially methylated regions (aDMRs) to those that mapped within 2 kb window of the identified genes (aDMGs) of the King penguin genome, excluding long-noncoding RNA aDMGs, resulting in 317 genes. We extracted the Illumina 450K probes mapping to human orthologs for these aDMGs, capturing 302 distinct Penguin aDMGs (N_CpG_=9,939).

We used randomized downsampling to infer the effect of sample size on the proportion of significantly age-associated CpGs. We randomly sampled our human data (N_Person_=1,501, N_Family_=761) to include 144 unrelated people, and ran our EWAS model without cell-type adjustment following the same pipeline as described above (excluding familial clustering). This process was repeated 100 times. The results across these 100 iterations were summarised to report the proportion of significant probes (FDR≤0.05) under a comparable data set as our penguin EWAS.

### PenguinAge development and application in humans

Using the above-described FTC data, we randomly selected one person from each family to compile a dataset of unrelated individuals for training the epigenetic clock for humans (N=761). The final dataset was randomly partitioned into a 70% training set and a 30% testing set. We restricted the methylation dataset to include CpG sites mapping to orthologues of the King penguin aDMGs (N=398), resulting in 302 genes (N_CpG_=9,939). An Elastic Net regression model was trained against age using the glmnetUtils package version 1.1.9 in R, with hyperparameters *α* and *λ* optimized via 10-fold cross-validation on the training set. The final model was built using the parameters that resulted in minimal prediction error (*λ*_min_ and corresponding *α*).

We applied the trained PenguinAge clock on the testing data along with other published epigenetic clocks, namely Horvath^22^, Hannum^64^, PhenoAge^65^ and GrimAge2^66^. Epigenetic ages of the above algorithms were calculated in R using the scripts provided in the corresponding publications. For each clock, epigenetic age acceleration (EAA) was determined as the residuals of epigenetic age linearly regressed against chronological age. Intrinsic EAA (iEAA) was calculated following Zhang et al. 2024^11^, by regressing epigenetic age against chronological age and relative proportions of 11 cell types (Bas, Neu, Bnv, CD4mem, CD4nv, CD8mem, CD8nv, Eos, NK, Mono and Treg).

To assess the associations between immune cell abundance and both EAA and iEAA measures, all cell-type fractions and age acceleration metrics were standardized using z-score transformation. Multivariate linear regression models were constructed for each combination of clock, metric, and cell type. The standardized EAA/iEAA was modeled as a function of individual cell-type proportion, adjusting for chronological age, squared chronological age (age^2^) and sex. Cell-type coefficient p-values were FDR adjusted to account for multiple testing across metrics, clocks and cell-types.

To evaluate whether PenguinAge is more robust to variation in blood cell composition than any clock created from conserved orthologues, we generated 1,000 clocks based on random gene sets. In each iteration, 409 genes were sampled from the set of King penguin orthologs that had at least one CpG site within 15kb up- and 2kb downstream (N_Gene_=16,465). For each randomly selected set of 409 genes, their corresponding human orthologue genes and CpG probes were identified, and an elastic-net clock was trained using the same training and test sets as used for PenguinAge. EAA and iEAA were calculated, and associations with estimated blood cell-type proportions were assessed as described above. For each cell type, we calculated the proportion of random clocks with weaker absolute standardized regression coefficients than PenguinAge. We additionally compared the mean absolute standardized regression coefficient across all cell-types between PenguinAge and the 1,000 random clocks as an overall measure of cell-type dependency.

## Acknowledgements

This study was funded and supported by the Research Council of Finland (grants #331320 and #354649 to RC), by the French Polar Institute Paul-Emile Victor (IPEV Project 137-ANTAVIA-POLAROBS, PI CLB), by the Centre National de la Recherche Scientifique (CNRS) through the Zone Atelier Antarctique et Terres Australes (ZATA), and by Zoo Zurich. The contribution of the Centre Scientifique de Monaco (CSM) to this work was supported by its institutional funding provided by the Government of the Principality of Monaco. This work was supported by the DFG Research Infrastructure NGS_CC (project 407495230) as part of the Next Generation Sequencing Competence Network (project 423957469). Sequencing was carried out at the Competence Centre for Genomic Analysis (Kiel). We especially acknowledge the key contribution of the Finnish IT Center for Science (CSC) for access to computational resources necessary to the realisation of this work. We also thank Prof. Jaakko Kaprio for providing data for humans used in this study. We are deeply grateful to the Zoo Zurich team, as well as to all the wintering and summering members of IPEV Project 137 and all the other colleagues and students within the P137 team, who participated in the long-term monitoring and sample collection since 2000. We also sincerely thank the IPEV logistics teams for their important and continued support in the field. This study is part of and supported by the long-term Studies in Ecology and Evolution (SEE-Life) program of the CNRS.

The estimation of cell type proportions in the human data was enabled by the use of the Software package developed at Dartmouth College, which software is subject to the licensing terms made available by Dartmouth Technology Transfer and which software is provided “AS IS” with no warranties whatsoever.

## Author contribution

The study was designed by RC, BSM, FANF and MHuk. Sample collection was done by RC, CLB, FANF, LD, MHof, GB and EP (penguins) and by MO (humans). Laboratory work was performed by RC, FANF, SZ, SW and LD. Analysis was designed and conducted by FANF, TZ and RC (penguins) and MHuk, RC and MO (humans). Manuscript elaboration was done by FANF, MHuk, BSM and RC. Manuscript revision was done by all authors. All authors read and approved the final manuscript.

## Supplementary Information is available for this paper

Requests for materials should be addressed to Robin Cristofari

