## Supplementary Figures for "An epigenetic palimpsest: the King penguin’s blood methylome reveals conserved hematopoietic ageing"

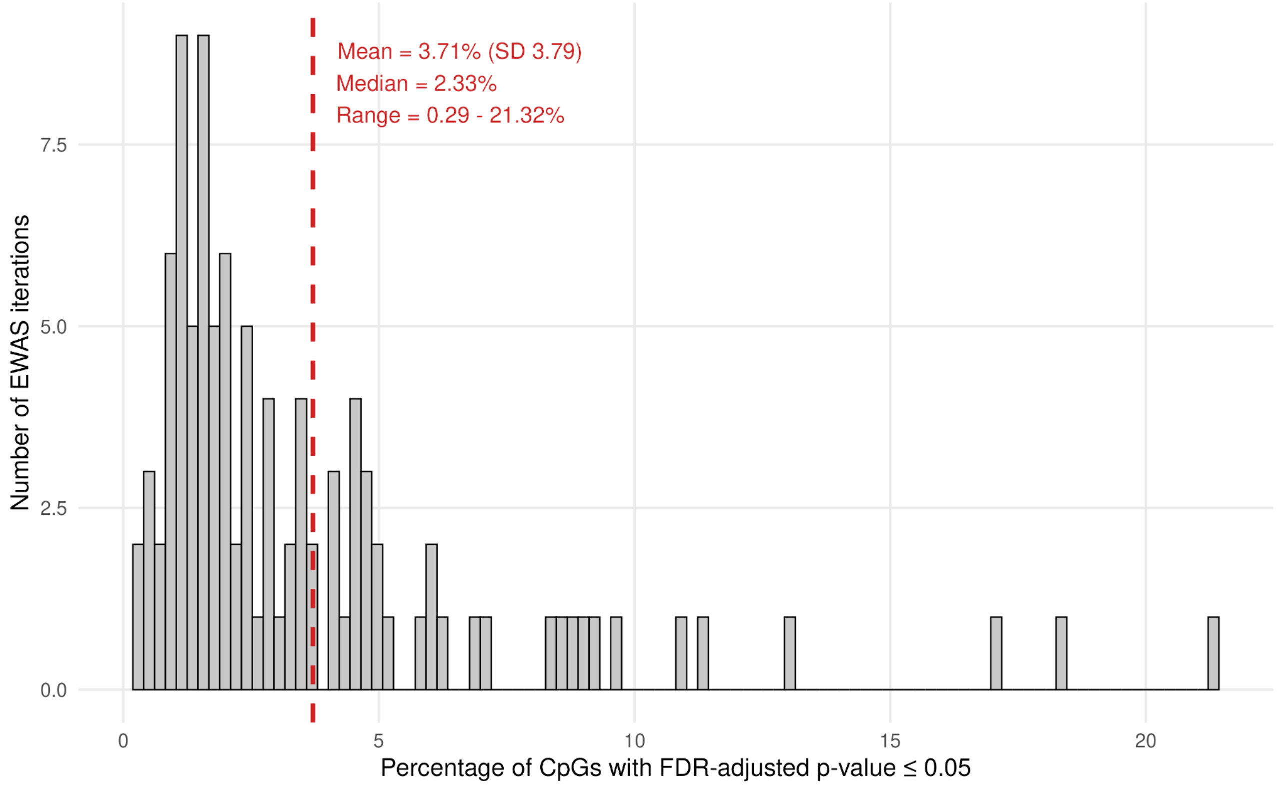


**Suppl. Fig. 1 | Distribution of the proportion of FDR-significant CpGs across 100 downsampled EWAS analyses.** Each downsampled replicate was performed in randomly selected subsets of unrelated individuals ($N_{Person}$=114, $N_{Family}$=114), and included the same probes ($N_{CpG}$=390,304) and covariates (BMI, smoking, alcohol, sex, technical factors) as our main analyses (excluding family ID). For each iteration, the percentage of tested CpGs with an FDR-adjusted p-value ≤ 0.05 was calculated. The dashed red line indicates the mean percentage across iterations.

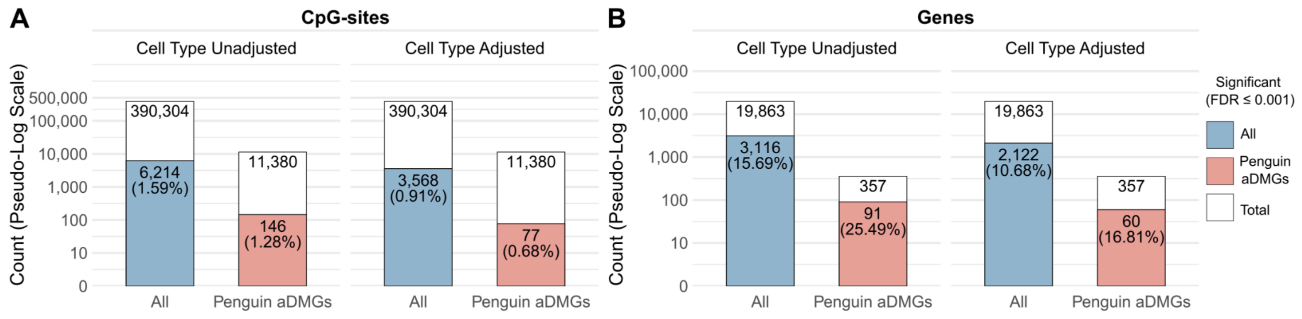


**Suppl. Fig. 2 | Epigenome-wide age association and epigenetic clock dynamics in humans. (A–B)** CpG and gene landscapes. Total counts and proportions of significant CpGs **(A)** and genes **(B)** evaluated with and without cell-type adjustment. Covariates for both models included BMI, smoking, alcohol, sex, technical factors, and family ID ($N_{Person}$=1,501. $N_{Family}$=761). Within nested bars, total features are white with significant fractions in blue; penguin aDMG orthologs are shown separately with significant fractions in red.

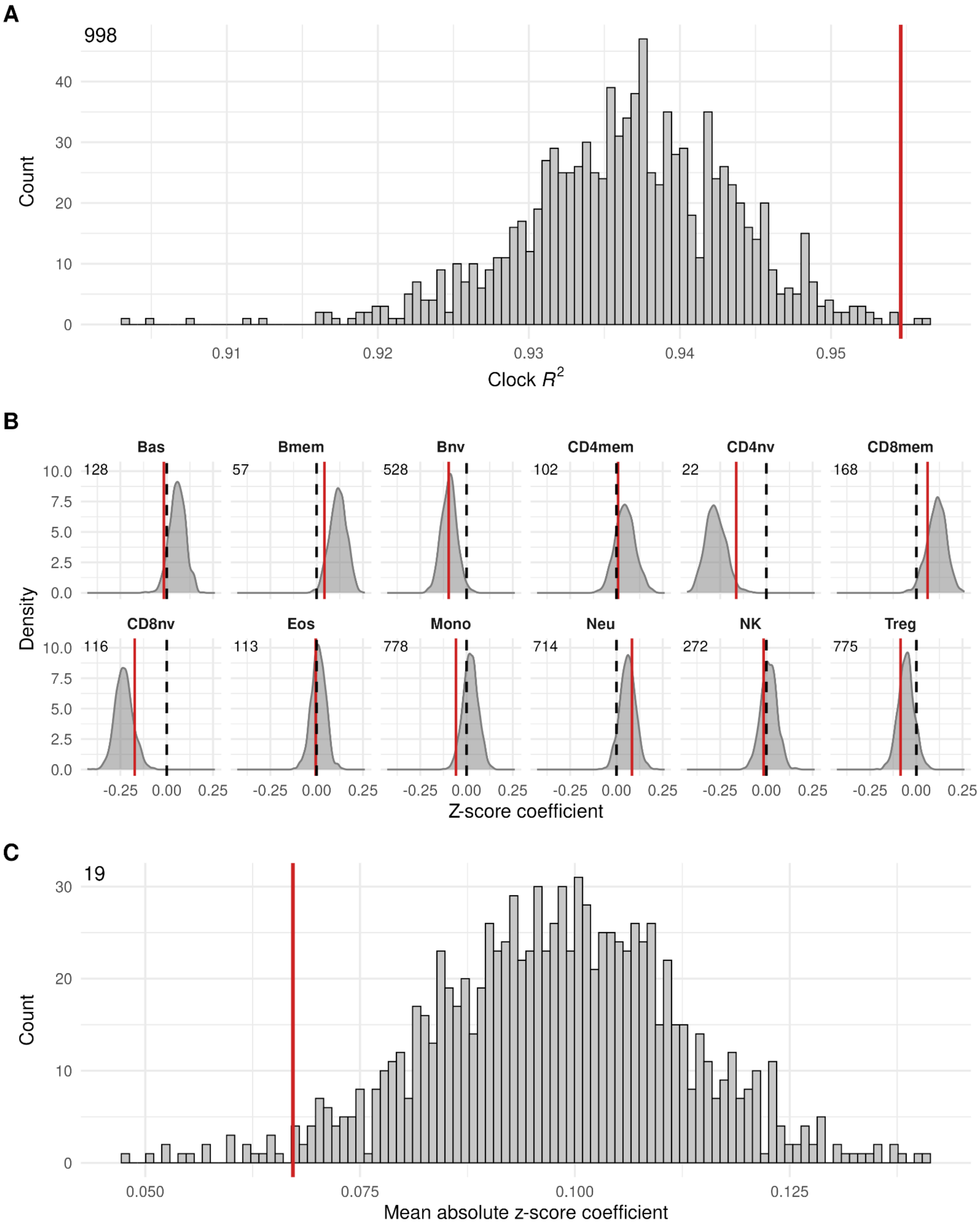


**Suppl. Fig. 3 | PenguinAge comparison with 1,000 random orthologue-based clocks.** Comparison of 1,000 clocks generated from human orthologs of randomly selected 409 King penguin genes. The clocks were evaluated in the same independent test set as used for PenguinAge (N = 229 unrelated individuals). Red lines denote the corresponding PenguinAge parameters, and upper left annotations indicate the number of clocks with smaller absolute estimates than PenguinAge. **(A)** Clock $R^2$ values between predicted and observed age. **(B)**Distributions of standardized regression coefficients relating EAA to 12 blood cell-type proportions. Dashed black lines indicate regression coefficient = 0. **(C)** Distribution of the mean absolute coefficient across all cell types for each random clock.
